# *Capitella teleta* gets left out: Possible evolutionary shift causes loss of left tissues rather than increased neural tissue from dominant-negative BMPR1

**DOI:** 10.1101/2023.10.04.560756

**Authors:** Nicole B. Webster, Néva P. Meyer

## Abstract

**Background:** The evolution of centralized nervous systems (CNSs) a fascinating and complex topic; further work is needed to understand the genetic and developmental homology between organisms with a CNS. Research into a limited number of species suggests that CNSs may be homologous across Bilateria. This hypothesis is based in part on similar functions of BMP signaling in establishing fates along the dorsal-ventral (D-V) axis including limiting neural specification to one region of ectoderm. From an evolutionary-developmental perspective, the best way to understand a system is to explore it in a wide range of organisms to create a full picture.

**Methods:** Here we expand our understanding of BMP signaling in Spiralia, the third major clade of bilaterians, by examining phenotypes after expression of a dominant-negative BMP Receptor 1 and after knock-out of the putative BMP antagonist Chordin-like using CRISPR/Cas9 gene editing in the annelid *Capitella teleta* (Pleistoannelida).

**Results:** Ectopic expression of the dominant-negative Cte-BMPR1 did not increase CNS tissue or alter overall D-V axis formation in the trunk. Instead, we observed a unique asymmetric phenotype: a distinct loss of left tissues including the left eye, brain, foregut, and trunk mesoderm. Adding ectopic BMP4 early during cleavage stages reversed the dominant-negative Cte-BMPR1 phenotype, leading to a similar loss or reduction of right tissues instead. Surprisingly, a similar asymmetric loss of left tissues was evident from CRISPR knock-out of *Cte-Chordin-like* but concentrated in the trunk rather than the episphere.

**Conclusions:** We further solidify the hypothesis that the function of BMP signaling during establishment of the D-V axis and CNS is fundamentally different in at least Pleistoannelida, possibly in Spiralia, and is not required for nervous system delimitation in this group. Our results support hypotheses of either multiple evolutionary origins of CNSs across Bilateria or divergence in the molecular mechanisms of CNS specification and D-V axis formation in annelids.

## 1. Background

The nervous system is an important animal innovation whose origins are poorly understood, especially the evolution of centralized nervous systems (CNSs). Many questions remain about the evolutionary and developmental processes that have allowed for the great diversity of extant nervous systems. Only after we have a full picture of nervous system diversity and how nervous systems develop can we start to answer big evolutionary questions: Have CNSs evolved repeatedly? What parts of CNSs may be homologous? What can that tell us about the evolution of complex systems in general? Are gene networks be repeatedly co-opted to form analogous systems?

In many bilaterian animals with a CNS, neural tissue is localized along the dorsal-ventral (D-V) axis, e.g., vertebrates have a dorsal neural tube, whereas arthropods and annelids have a ventral nerve cord. A key part of D-V axis and CNS formation in some vertebrates and ecdysozoans are the Bone Morphogenetic proteins (BMPs), which are within the Transforming Growth Factor β (TGF-β) superfamily. A gradient of BMP signaling helps establish fates along the D-V axis including limiting where neuroectoderm forms (1). Later in development, a BMP gradient patterns neural subtypes along the D-V axis of the CNS itself (2). As with other members of the TGF-β superfamily, BMPs are secreted ligands that dimerize and bind to a tetrameric, extracellular receptor complex made up of two type I and two type II receptors (3). The type II receptor phosphorylates the type I receptor once the ligand is bound, causing the type I receptor to phosphorylate a transcription factor, Suppressor of Mothers against Decapentaplegic (SMAD), which then binds a co-SMAD and translocates into the nucleus to affect gene transcription. Secreted BMP inhibitors such as Chordin/Short gastrulation (Chd/Sog) are proposed to be the key regulators of the BMP signaling gradient (4). Thus, BMP’s anti-neural function is intrinsically linked to its organizing function in D-V axis formation (2).

One prevailing hypothesis suggests that a CNS evolved once, near the base of Bilateria, such that the brains and nerve cords of all bilaterians are homologous (5–7). This hypothesis is intertwined with the idea of axis inversion, where the CNS and D-V axis became inverted in the last common ancestor of deuterostomes, which could explain why BMP signaling is anti-neural in both groups, forming a dorsal nerve cord in vertebrates and a ventral nerve cord in insects. Recent work has refuted this hypothesis, at least partially because the function of BMP signaling during D-V axis and CNS formation differs between spiralians and the rest of Bilateria (8–12). Additionally, work outside traditional lab species in other lineages shows that within both deuterostomes (Enteropneusta: (13);) and ecdysozoans (Nematoda, Onychophora (14), the role of BMP signaling in D-V axis and neural specification is not as simple nor as conserved among animals.

To understand the evolution of key animal features, it is essential that we examine a broad range of animals. One key aspect of understanding a phenotype is to know the underlying genetic and developmental mechanisms responsible for its formation. A great deal is known about these genetic and developmental mechanisms in a few specific vertebrate and ecdysozoan organisms (e.g., frog, mouse, chick, fly, and *C. elegans*) (15–18). Yet little is understood about the genetic mechanisms underlying development in the third major clade of bilaterians, Spiralia (Lophotrophozoa *sensu lato*) (19–21). Beyond increasing taxonomic diversity to understand the evolution and development of phenotypes, Spiralia includes a great diversity of morphology and life histories within annelids, molluscs, brachiopods, rotifers, and platyhelminths (22). Furthermore, spiralian development is interesting in its own right: spiralian embryos cleave using a fascinating stereotypic cleavage pattern, spiral cleavage, which is considered ancestral for the clade (23,24). Each cell (blastomere) contributes to a stereotypical set of fates, some of which are labile, that arise through a combination of autonomous and conditional signaling (25). This mode of development is superficially similar to early ascidian development, giving us the opportunity to compare convergent evolution of genetic and cellular interactions in a constrained, possibly contact-dependent, mode of early development. Furthermore, spiral cleavage enables us to compare development of tissues arising from homologous blastomeres across spiralian taxa with very different body plans (e.g., a bivalve versus an annelid), giving us powerful tools to trace evolution of developmental mechanisms. Overall, there a number of aspects of spiralians that deserve added study, both in their own right, but also to diversify our comparative studies.

Within spiralians, the role of BMP signaling during CNS fate specification and D-V axis formation has been difficult to pinpoint (11,26). For example, in the mollusk *Ilyanassa* (∼*Tritia* (27)) *obsoleta,* BMP signaling appears to play a role in D-V organization, where a loss of BMP signaling caused a loss of the D-V axis, but not in repressing neural tissue formation (28). Instead, ectopic BMP caused ectopic eye and brain formation. In contrast, in the mollusc *Crepidula fornicata,* ectopic BMP caused a partial loss of the head (episphere) but a normal trunk (26). A more complex study in *Lottia goshimai* showed that perturbations of BMP or Chordin affected both eye number and D-V axis organization, but how CNS tissue was affected is unclear (29). In the annelid *Platynereis dumerilii,* ectopic BMP shifted the D-V boundaries of gene expression in the neuroectoderm, but did not shift neuroectodermal boundaries as assayed by expression of the pan-neuronal gene *Pdu-elav* (7). In the leech *Helobdella*, gain and loss of BMP signaling affected the D-V identity of the o and p ectodermal bandlets in the trunk but not in the rostral segments; no effect was reported for the neural (n) bandlet (30,31). In the annelid *Capitella teleta,* we previously showed that ectopic BMP does not reduce neural tissue or affect D-V axis formation (11). Instead of BMP, Activin/Nodal organizes the D-V axis in *C. teleta* (8,9,32). Further complicating this story, our lab has demonstrated a decoupling between neural specification and D-V axis formation and between neural specification in the head versus trunk in *C. teleta* (33). Overall, these diverse results raise questions about the conservation of BMP signaling during D-V axis and CNS formation in Spiralia and compared to other bilaterians.

Our lack of consensus of how BMP signaling functions in spiralians is at least partially due to a lack of functional studies in spiralians and to differences in methodology across studies. In *C. teleta,* the spiralian we are studying, there are two BMP ligands, Cte-BMP2/4 and Cte-BMP5–8, and two BMP receptors, a type 1, Cte-BMP Receptor 1 (BMPR1=Alk3/6 Activin receptor-like kinase]) and a type 2, Cte-BMP Receptor 2 (BMPR2) (34). Based on models of BMP signaling in vertebrates and insects, both ligands are thought to signal to the nucleus using the phosphorylated receptor-regulated SMAD, SMAD1/5/8, and a co-SMAD, SMAD4, although ActivinR1 (ALK1/2) may also bind BMP5–8 and transmit through the other rSMAD, SMAD2/3 (3). A key antagonist in the system, Chordin, normally regulates the BMP gradient, however Chordin has been lost in most annelids (Pleistoannelida) but Chordin-like may play a similar role (34,35).

We previously published work showing that incubating *C. teleta* embryos in ectopic BMP does not disrupt overall D-V axis formation or reduce the amount of CNS tissue formed (11). Here we examine the effect of altering BMP signaling in *C. teleta* using a dominant-negative Cte-BMPR1 and knock-down of Cte-Chordin-like via CRISPR/Cas9 gene editing. We truncated the kinase domain of BMPR1, creating a dominant-negative BMP receptor (BMPR1ΔK) that decreased downstream phosphorylation of SMAD 1/5/8 but did not increase CNS tissue or alter overall D-V axis formation in the trunk. Instead, BMPR1ΔK injection resulted in a unique asymmetric phenotype: a distinct loss of left tissues including the left eye, brain, foregut, and trunk mesoderm. A similar asymmetric loss of tissue was evident from CRISPR knock-out of Cte-Chordin-like. Overall, we show added symmetry related functions of BMP signaling in spiralians, but further reinforce no neural specification role.

## 2. Material and methods

### 2.1. Animal care and embryo collection

Adults of *Capitella teleta* Blake et al. (36) were cultured in glass finger bowls with 32–34 ppt artificial sea water (ASW; Instant Ocean Sea Salt in Hydro Picopure-filtered tap water) at 19 °C and fed with sieved mud collected from the local coastline (37–39). In order to collect embryos of the correct stage (st.), mating dishes were generated by separating males and females for 3–5 days at 19 °C and then either 1) combining males and females for 5–16 h in the dark at 19 °C or 2) exposing males and females for 6+ h to light at room temperature (RT, ∼21°C) and then combining them for 5 h at RT (32). Embryos and larvae, except where otherwise noted, were raised in ASW with 50 μg/mL penicillin and 60 μg/mL streptomycin (ASW+PS) at RT. ASW+PS was changed once or twice daily.

### 2.2. Isolation of *C. teleta BMP receptor 1*

Total RNA was extracted from mixed stage 1–9 embryos and larvae using the RNA Trizol extraction protocol (Molecular Research Center, Inc.) or the RNeasy Mini Kit (Qiagen cat. 74104) paired with the QIAshredder columns (Qiagen 79656). Reverse transcription reactions were conducted using the SMARTer RACE kit (Clontech 634859) or High-capacity cDNA Reverse Transcription kit (Applied Biosciences 4368814). Only one BMP Receptor 1 (Cte-BMPR1;) homolog has been identified in the *C. teleta* genome (34). A 1533 bp fragment of Cte-BMPR1 (JGI PID111904) encoding nearly the entire coding sequence (but lacking the last 54 bp at the end of the 3’ UTR) plus 25 bp of 5’-UTR was amplified by PCR using 3 sets of overlapping gene-specific primers, followed by a nested PCR using the SMARTer RACE kit following the manufacturer’s protocol. The sequence we isolated was confirmed via BLAST (40) with 67% protein sequence identity compared to *Platynereis dumerilii* (CAE76647.1) and 66% with *Lamellibrachi satsuma* (KAI0231297.1). The retained domains include the BMP binding domain, transmembrane domain, and GS domain.

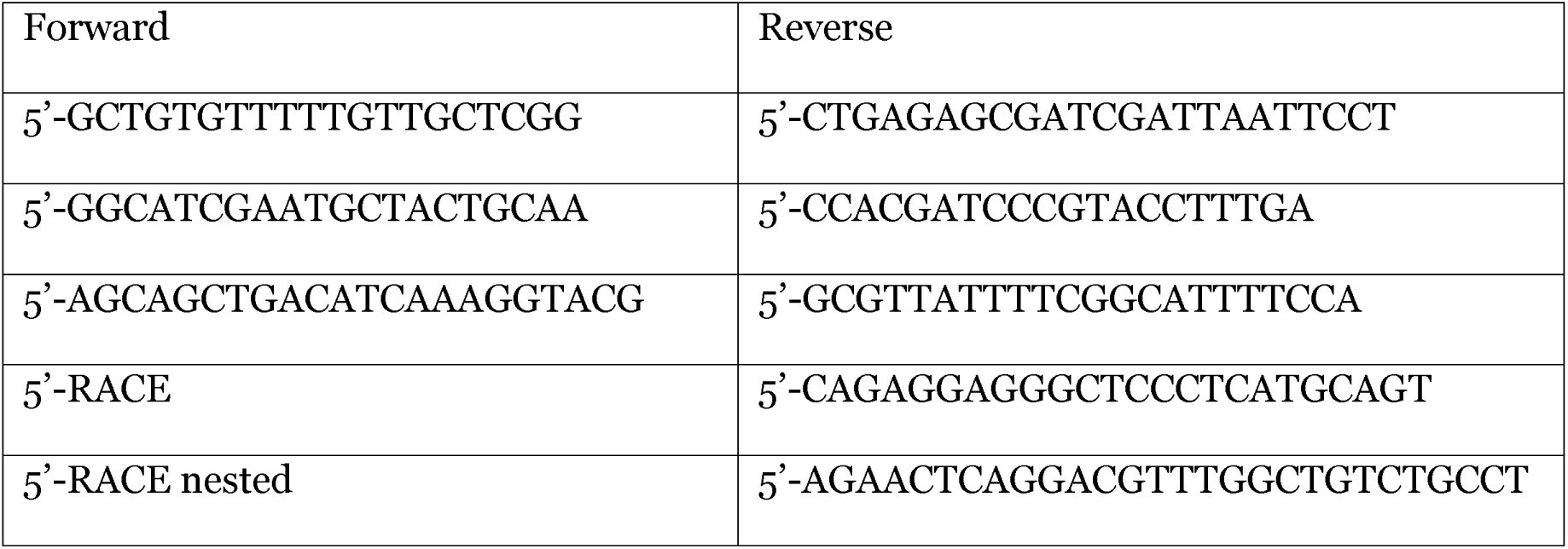

### 2.3 BMPR1ΔK construction

Cte-BMPR1 domains were determined by aligning protein sequences with previously published BMPR1 sequences (*Drosophila melanogaster*, AAA61947.1, *Xenopus laevis*, BAA22438.1, *Platynereis dumerilii* CAE76647.1, *Helobdella sp. Austin* JN091774.1). The dominant-negative BMPR1ΔK was designed by truncating Cte-BMPR1 at aa 221 to remove the kinase domain, following previously-designed dominant-negative BMPR1 constructs (31,41,42); see Supplemental Fig. 1. The designed dominant-negative sequence (aa 1–220) was synthesized (Eurofins) for use in the Gateway system (Invitrogen) and ligated into an entry vector using the pENTR/D-TOPO Cloning Kit with One Shot™ TOP10 Chemically Competent *E. coli* (Invitrogen K2400-20). Following Gateway manufacturing protocols, we constructed a fusion protein with Cte-BMPR1ΔK and mVenus using pSPE3-RfA-Venus (43) and LR Clonase II (Invitrogen 11791-020), and the final expression vector, pSPE3 Cte-BMPR1ΔK::mVenus, was verified by sequencing. mRNA was then transcribed using the mMESSAGE mMACHINE™ T3 Transcription kit (Invitrogen AM1348), a polyA tail was added with the Poly(A) Tailing kit (Invitrogen AM1350), and mRNA was purified with the MEGAclear kit (Invitrogen AM1908) and concentrated with an ammonium acetate precipitation. mRNA was resuspended in RNAse-free water (Invitrogen AM1908), and a Nanodrop One (Thermo Scientific) was used to determine concentration. BMPR1ΔK::mVenus mRNA was then aliquoted for single-use and stored at -80 °C following Layden et al. (44) and Özpolat et al. (45).

### 2.4 CRIPSR/Cas9

Two 20 bp sgRNA target sequences (sgRNA175: AGTGCCGCAAGACTCTTGTG; sgRNA264: CCACGGGAGTCGTGTATCCA) and amplification primers were designed using CRISPOR (46) for NGG PAM sequences near the 5’ end of *Cte-chordin-like* (*Cte-chd-l;* JGI PID224618)(34,47) with minimal off target complementarity with the *C. teleta* genome. Complete sgDNA templates were assembled and amplified using the T7 promoter (Phusion High Fidelity, NEB E0553). sgRNA was then transcribed (MEGAshortscript T7, Invitrogen AM1354), cleaned (RNA Clean and Concentrator, Zymo Research R1013) and aliquoted to 1 µg/µL in RNAse-free water at -80 °C for storage (concentration was determined using a Qubit 3 Fluorometer, Invitrogen).

Verification of *in vitro* cutting was confirmed by adding 250 ng of each sgRNA and 500 ng of Cas9 protein (PNA Bio cat. CP01-200) to 250 ng of PCR-amplified *Cte-chd-l* and incubating at RT for 1 h. The sample was then run on a 1% agarose gel. The expected cut site was amplified via PCR using *Cte-chd-l* specific primers (175L: CGAGAGGACGACAACCAGAG; 175R: TTGTGCGTTTCCTGCGAAAG). Verification of i*n vivo* Cas9 cutting of *chd-l* following Neal *et al.* (48). Briefly, the genomes of individual stage 6 larvae were extracted after Cas9/sgRNA injection as zygotes and sent for sequencing using ­*Cte-chd-l* specific primers. Of five embryos that were sequenced, four showed evidence of cleavage via CRISPR.

### 2.5 Microinjection

Prior to injection, the outer egg envelope of zygotes or early cleavage-stage embryos was permeabilized for 30 sec using a freshly mixed, 1:1 solution of 1 M sucrose and 0.25 M sodium citrate (individual solutions were prepared the previous day, stored at 4 °C and then warmed to RT before use). Egg envelope permeabilization was followed by three rinses with ASW+PS. DiI injections into cleavage-stage embryos were conducted following Meyer et al. (49). mRNA and sgRNAs/Cas9 were injected into zygotes. In *C. teleta*, the time to first cleavage after fertilization has not been carefully determined but appears to be ∼4–6 h at 19 °C. Because fertilizations are likely internal in *C. teleta* (50), the precise timing of fertilization for the collected zygotes was unknown. In general, zygotes started cleaving to two cells anywhere from a few minutes to a few hours after injection. mRNA injections were performed with beveled (Sutter BV-10) Quartz needles (Sutter Instrument Co., Novata, CA, USA), where needles were pre-warmed to 55 °C before being backfilled. The injectant was mixed with 5x Rhodamine-Dextran (30 mg/mL Dextran Tetramethylrhodamine 10k MW Invitrogen D1868) as a tracer and a 5x injection buffer (10 mM HEPES pH 7.0; 75 mM KCl) (51) in RNAse-free water to a final concentration of 1x for both. BMPR1ΔK mRNA concentrations ranging from 480 ng/µL to 1.6 µg/µL were injected with no changes in resulting phenotypes. For *Cte-chd-l* CRISPR, needles were backfilled with 125 ng/µL of each sgRNA and 2 µg/µL Cas9 protein, rested for 10 min at RT, then stored at 4 °C for up to 5 days. Some animals were mounted in ASW+PS for live imaging of mVenus expression (AxioImager M2 microscope (Zeiss); coverslips were sealed with vacuum grease to reduce evaporation. Both injected animals and uninjected controls from the same brood were raised at RT in ASW+PS until stage 6, and then animals were fixed and labeled for phenotypic scoring. An experiment was not scored unless 90% of the uninjected animals were healthy.

### 2.6 Incubation in BMP protein

To understand the interaction between BMPR1ΔK and BMP protein, BMPR1ΔK-injected animals (1 biological replicate) and uninjected control animals were incubated in 250 ng/mL BMP4 protein in ASW+PS for 12h starting at either at the 8-cell stage (first-quartet of micromeres or “1q”) or just after birth of micromere 4d (∼64-cell stage or “4q”). Stock recombinant zebrafish BMP4 protein (R&D Systems 1128-BM-010) was reconstituted to 20 µg/mL in 0.1% bovine serum albumin (BSA) and 4 mM HCl in Picopure water, aliquoted and stored at -80°C (11). Animals were raised until stage 6 in ASW+PS and then fixed to assess the resulting phenotypes.

### 2.7 Whole mount in situ hybridization

Whole mount in situ hybridization (WMISH) was conducted as described previously (52). Briefly, all WMISH fixations were done in 4% paraformaldehyde (PFA, stock 32% PFA ampules from Electron Microscopy Sciences, cat. 15714) in ASW for 6 h–overnight at 4 °C. After fixation, animals were serially dehydrated in methanol and stored at -20 °C. Animals were hybridized for a minimum of 72 h at 65 °C with 1 ng/µl of each probe. Spatiotemporal RNA localization was observed using an NBT/BCIP color reaction. The color reaction was stopped using 3 washes of PBS with 0.1% Tween-20. After WMISH, animals were labeled with Hoechst and anti-acetylated-Tubulin (details below), cleared in 80% glycerol in PBS and mounted on slides for DIC and fluorescent imaging.

### 2.8 Fixation, staining and antibody labeling in larvae

Prior to fixation, the egg envelope of embryos was permeabilized for 3 min using a freshly mixed 1:1 solution of 1 M sucrose and 0.25 M sodium citrate. In the case of larvae, they were relaxed in 1:1 ASW:0.37 M MgCl_2_ for 5–10 min before fixation. Immunolabeling was carried out as in Meyer et al. (2015). Animals were fixed for 30 min with 4% PFA in ASW at RT, rinsed with PBT (PBS + 0.1% Triton-X 100), blocked in 5 or 10% heat-inactivated goat serum in PBT (block) and incubated in primary antibody in block overnight at 4 °C. Secondary antibodies in block were incubated overnight at 4 °C, then animals were thoroughly washed with PBT and cleared and mounted in SlowFade Gold (Life Technologies, cat. S36936) for confocal laser scanning microscopy or in 80% glycerol in PBS for all other types of microscopy. All washes and exchanges were done in RainX-coated (RainX) glass spot dishes. Primary antibodies used were as follows: 1:800 rabbit anti-serotonin (5HT; Sigma-Aldrich, cat. S5545), 1:20 mouse anti-Futsch (clone 22C10, Developmental Studies Hybridoma Bank), 1:800 mouse anti-acetylated-Tubulin (ac-Tub; clone 6-11B-1, Sigma, cat. T6793), and 1:400 rabbit anti-phosphorylated-SMAD1/5/8 (pSMAD1/5/8; clone 41D10, Cell Signaling Technologies). Secondary antibodies used were as follows: 1:2000 goat anti-mouse F(ab’)2 Alexa488 (Sigma-Aldrich, cat. F8521) and 1:1000 sheep anti-rabbit F(ab’)2 Cy3 (Sigma-Aldrich, cat. C2306). F-actin and DNA staining were performed by incubating the embryos and larvae in 1:100 BODIPY FL-Phallacidin (Life Technologies, cat. B607; stock concentration 200 Units/mL in methanol), 0.1 μg/mL Hoechst 33342 (Sigma-Aldrich, cat. B2261) along with the secondary antibodies.

### 2.9 pSMAD immunolabeling in cleavage-stage embryos

To detect levels of BMP signaling after injection, the amount of phosphorylated-SMAD1/5/8 in the nucleus was measured (1 biological replicate). Embryos were separated into four treatments: Uninjected embryos incubated for 1 h in ASW or BMP; BMPR1ΔK-injected embryos incubated 1 h in ASW or BMP; 250 ng/mL recombinant BMP4 was added at 4q (∼8-10h after injection) as previously described (11). Then the embryos were fixed for 15 min in 4% PFA at RT, and all other steps were carried out as above. Animals were labeled with 1:400 anti-phosphorylated-SMAD1/5/8 (clone 41D10, Cell Signaling Technologies), BODIPY FL-Phallacidin, and Hoechst 33342 as above.

All animals went through immunolabeling at the same time using the same protocol, and images were all taken on the same microscope (Confocal TCS SP5-X, Leica) using the same settings to control for differences in fluorescence. pSMAD1/5/8 levels were determined by averaging the fluorescence brightness of the anti-pSMAD1/5/8 antibody in the nucleus of the surface-most cells that were intact (not dividing) and did not appear distorted by the edge of the embryo. These 3–5 nuclei per animal were each measured 3 times with a newly drawn ROI on different days to reduce measurement bias and averaged in Leica Applications Suite X (Leica). Since embryos were imaged from different orientations, this meant that different cells were measured for each embryo.

### 2.10 Microscopy and figure preparation

Images were taken using DIC optics on an AxioImager M2 microscope (Zeiss) with an 18.0-megapixel EOS Rebel T2i digital camera (Canon) for WMISH animals or an AxioCam MRm rev.3 camera (Zeiss) with Zen Blue software (Zeiss) for antibody-labeled animals or live imaging of mVenus. DiI-labeled animals were imaged using a Zeiss Apotome.2 to produce optical sections. Animals for confocal laser scanning microscopy were imaged using a TCS SP5- X (Leica). DIC images taken at different focal planes were merged with Helicon focus 7 (Helicon). Different channels and z-stacks of fluorescent images were merged using Zen Blue (Zeiss). WHISH images were edited for contrast and brightness using Adobe Photoshop CC (Adobe). Figure panels were assembled with Adobe Illustrator CC (Adobe).

### 2.11 Statistics and analyses

Only elongated animals (ellipsoid body shape with all of the following: brain, VNC, prototroch and telotroch) were scored for phenotypes.

All statistics were performed in R/RStudio 1.2.5 (R Core team, 2014; RStudio Team, 2012), and all graphs were created using the ggplot2 package (Wickham, 2009) and polished with Adobe Illustrator CC (Adobe). Model testing was used determine the appropriate covariables to analyze in each ANOVA; the R package rcompanion (Mangiafico., 2015) was used, and the model with the lowest AIC was chosen. Tukey post-hoc analyses were used to determine the differences between treatments.

## 3. Results

### 3.1 BMPR1ΔK-injected animals expressed mVenus

To determine if BMPR1ΔK was expressed in embryos, we looked for the expression of the mVenus tag after injection using live imaging. mVenus fluorescence was first observable during early cleavage, 4–7 h after *BMPR1ΔK::mVenus* mRNA was injected into zygotes (n = 3 biological replicates), Fig. 1 shows mVenus 20 h after injection), and fluorescence was generally not detectable by 36 h (n = 8 biological replicates). However, mVenus fluorescence lasted much longer in two replicates, until st 5, which are larvae that just beginning to move by ciliary beating (∼4 days post injection). mVenus fluorescence was detectable in most cells in cleavage-stage embryos and was localized to cellular membranes, with additional punctate fluorescence surrounding the nucleus. This suggests a low degree of mosaicism and that the truncated receptor protein was properly localized to the membrane. While the intensity of mVenus fluorescence varied between replicates and individuals, injected animals showed similar phenotypes, even in animals with no *observable* mVenus fluorescence. This suggests that BMPR1ΔK protein is produced across a range of mRNA concentrations and is able to function similarly, even in embryos where mVenus fluorescence is not detectable.

**Figure 1.**
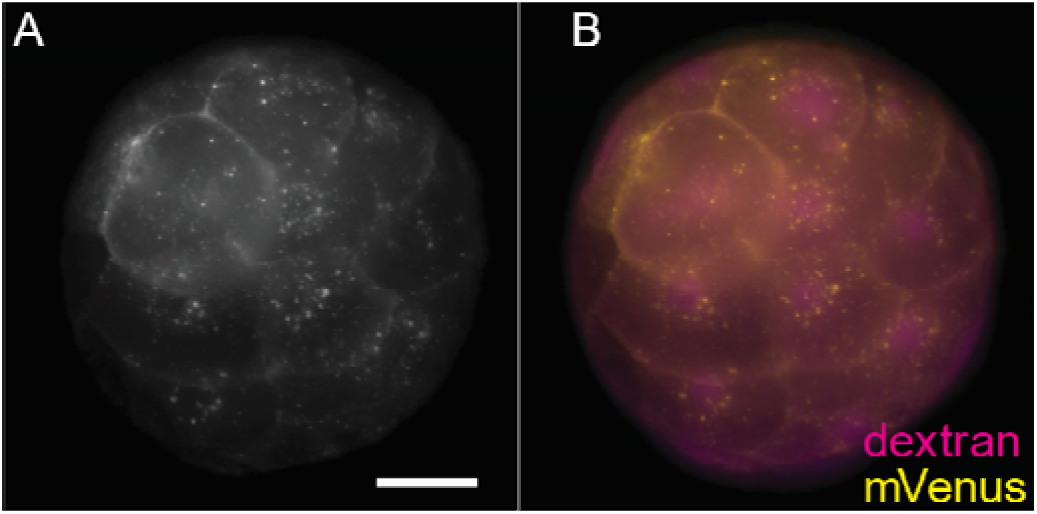
mVenus expression 20 hour after injection of 1.6 µg/µL mRNA encoding BMPR1ΔK::mVenus into a zygote. A. mVenus. B. merged image; mVenus (yellow) and Rhodamine-Dextran (magenta, tracer dye). Scale bar: 50 µm.

### 3.2 BMPR1ΔK can reduce nuclear pSMAD1/5/8

To determine if BMPR1ΔK affected downstream signaling of the BMP pathway, we assayed pSMAD1/5/8 expression in embryos after injection. We previously showed that ectopic BMP4 increased pSMAD1/5/8 expression in *C. teleta* (11), and this was used as a positive control. One biological replicate demonstrated the expected pSMAD1/5/8 response to BMPR1ΔK injection (Fig. 2). Uninjected controls and BMPR1ΔK-injected embryos (with or without ectopic BMP4) generally had low to non-detectable amounts of nuclear pSMAD1/5/8, whereas uninjected embryos with ectopic BMP4 showed a significantly higher level of nuclear pSMAD1/5/8 than either uninjected controls and BMPR1ΔK-injected embryos (ANOVA, F_treatment_ = 150.75, df = 3, p < 0.0001; Tukey HSD p < 0.0001).

**Figure 2.**
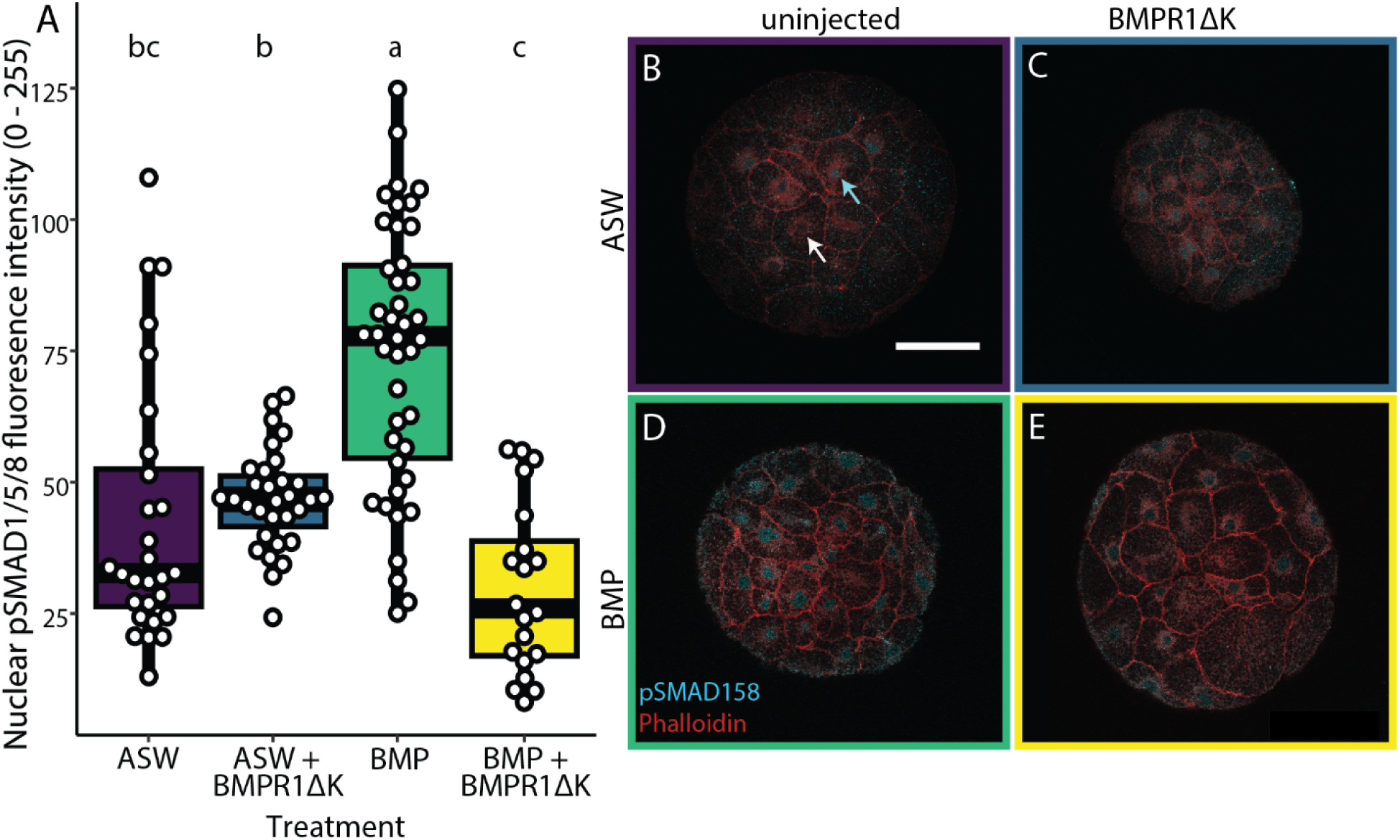
Nuclear pSMAD1/5/8 levels approximately 10 h after BMPR1ΔK injection. A. Boxplot showing the varying effect of BMPR1ΔK injection on the amount of nuclear pSMAD1/5/8 (measured as the relative brightness of labeling with an anti-pSMAD1/5/8 antibody in the nucleus); letters indicate significance groups; white dots indicate individual cells. B) Uninjected embryo in ASW, Arrows: cyan: higher nuclear pSMAD1/5/8; white: lower/no nuclear pSMAD1/5/8. C) BMPR1ΔK-injected embryo in ASW. D) Uninjected embryo with 1 h BMP4 pulse. E) BMPR1ΔK-injected embryo with 1 h BMP4 pulse. Cyan: pSMAD1/5/8; Magenta: Phalloidin. Scale bar: 0.5 µm.

In some uninjected controls, nuclear pSMAD1/5/8 levels varied between non-dividing cells; some nuclei appeared to have higher levels of pSMAD1/5/8 compared to their neighbors (cyan versus white arrows in Fig. 2). Interestingly, the BMPR1ΔK-injected embryos did not appear to have the same variation in nuclear pSMAD1/5/8 levels within an embryo, with or without added BMP4.

In summary, BMPR1ΔK injection was able to block an increase in nuclear pSMAD1/5/8 when ectopic BMP4 was added, suggesting that BMPR1ΔK protein is able to block activation of the BMP pathway in *C. teleta*.

### 3.3 BMPR1ΔK injection produced left-reduced, asymmetric features

The majority (55.6% ± 5.3 SE) of BMPR1ΔK-injected zygotes (n = 420, 13 replicates) did not elongate. Un-elongated embryos presented a broad range of features, but were generally more spherical than controls, and none had all of the following: brain, VNC, prototroch, telotroch (Fig. 3). There was a no significant effect of *BMPR1ΔK::mVenus* mRNA concentration on the proportion of elongated embryos, but injected embryos with added ectopic BMP4 did elongate significantly more often (82.4% ± 0.1 SE; n = 2; T-test, df = 12, t = 3.0, p < 0.006). Elongated animals were further scored for phenotypic changes relating to BMPR1ΔK-injection.

**Figure 3.**
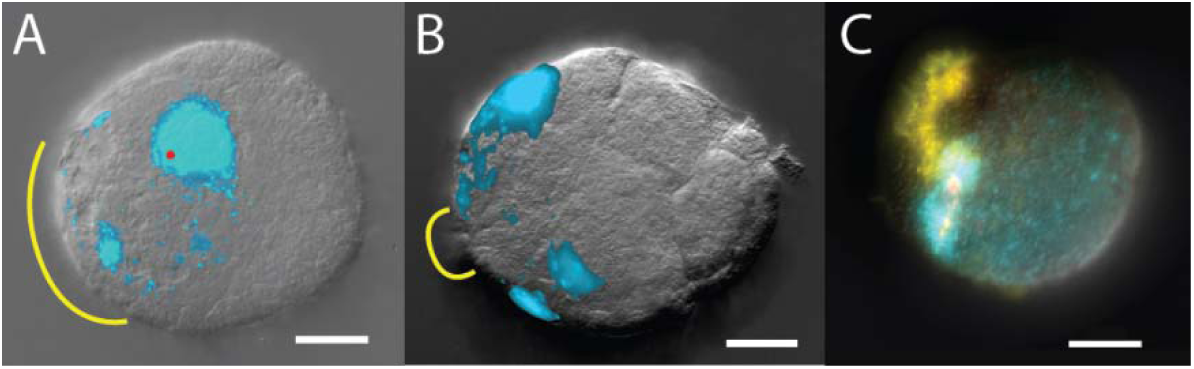
Representative unelongated larvae after BMPR1ΔK-injection. A) Anterior view of animal with 1 eye pigment cell (red), and 1 presumed brain lobe. B) larva (view unknown) with no clear features. C) Anterior view of animal with 1 eye sensory cell (red). Yellow: cilia; cyan: nuclei; red: larval eye; scale bars: 0.5 µm.

While most elongated BMPR1ΔK-injected animals were relatively normal at stage 6, the most common, non-wild-type phenotype was a general reduction or loss of tissue on the left side. This included the brain lobe and larval eye on the left side of the episphere, and the foregut, mesodermal band and muscle fibers on the left side of the trunk (Figs. 4, 5, 6). This trait asymmetry appears to be correlated; the asymmetry of brain lobes, foregut, or mesoderm tissue was not independent (pairwise Fisher’s exact tests, p < 1×10^-09^) such that animals lacking a left brain lobe were more likely to also lack their left mesoderm than chance. Furthermore, concentrations of *BMPR1ΔK::mVenus* mRNA ranging from 480 ng/µL to 1.6 µg/µL were injected with no significant changes in the proportion of resulting phenotypes (foregut, brain, and mesoderm asymmetry; t_paired_(2) = 2.4, p=0.14).

**Figure 4.**
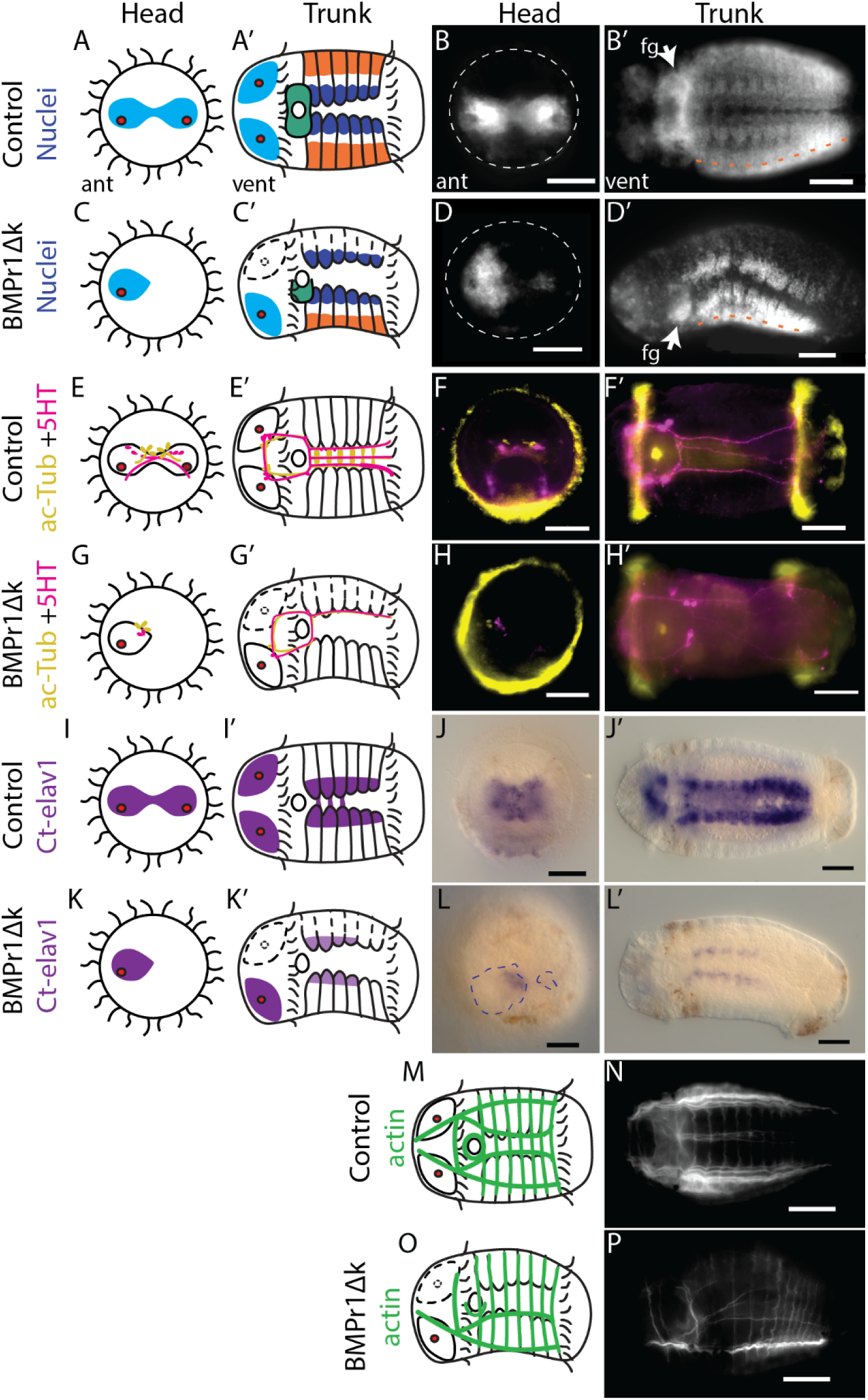
Diagrams and images of representative control and BMPR1ΔK-injected animals. A–D) Phenotypes visible via nuclear staining (Hoechst): brain lobes (cyan), foregut (green), hemiganglia of the ventral nerve cord (blue), and mesodermal bands (orange). White dashed lines: episphere outline; orange dashed line: division between mesoderm and non-neural ectoderm. E–H) Neurons and neurites (ac-Tub: yellow, 5HT: magenta). I–L) Post-mitotic neurons (anti-*Cte-elav1* ISH) Blue dashed lines: brain. M–P) Muscle fibers (Phalloidin). See text for numbers of animals. For D, H and L, the anterior and ventral views are from different animals to illustrate the generalized phenotype. Scale bars: 0.5 µm.

**Figure 5.**
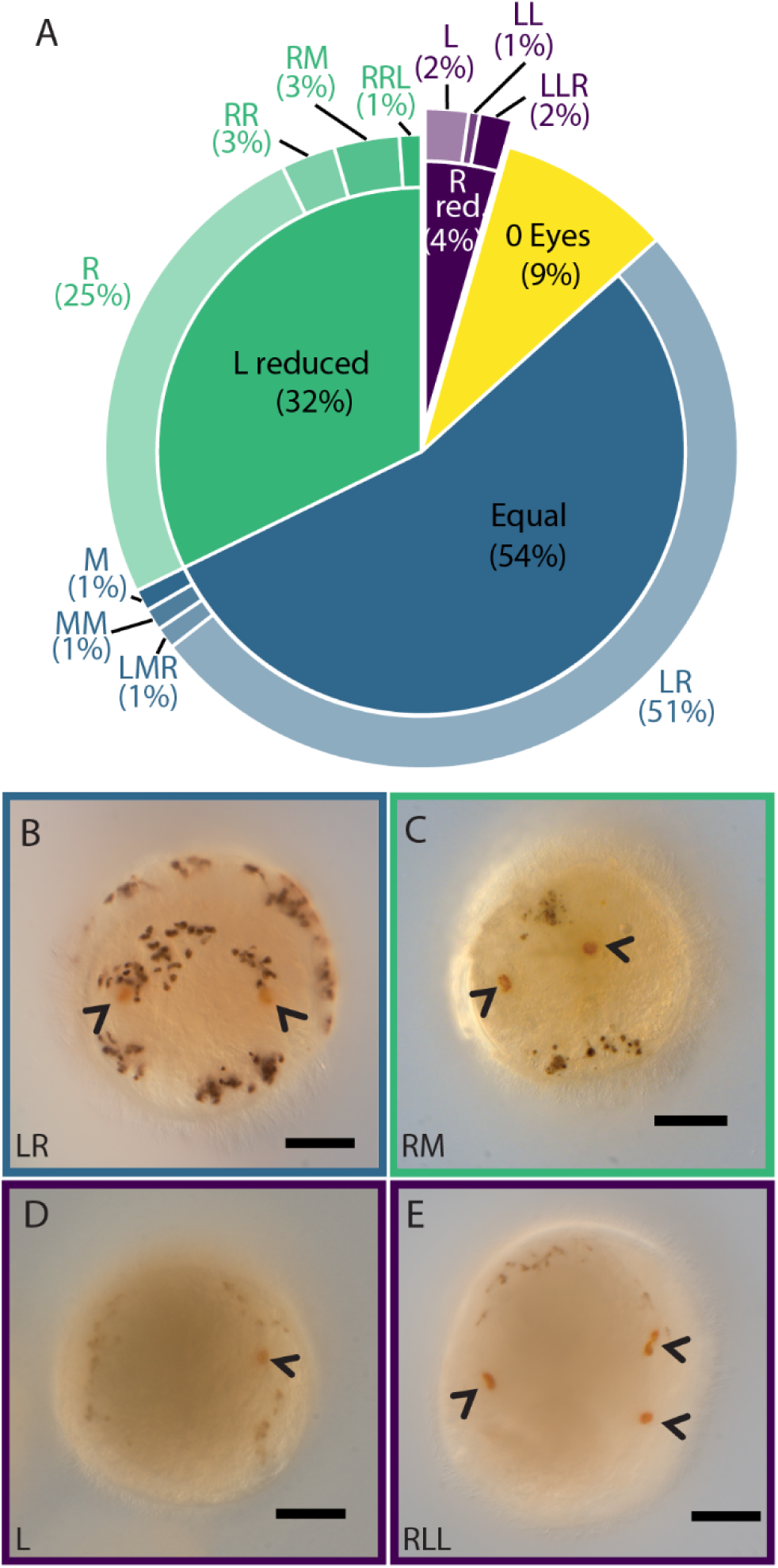
Eye formation after BMPR1ΔK-injection A) Pie chart showing the proportion of each larval eye pigment cell phenotypes: Equal (blue), right-biased (green) and left-biased patterns (purple) were further sub-divided based on the number of eye pigment cells in each position, e.g., RM indicates 1 right and 1 medial eye pigment cell; R, right; L, left; M, medial. B–D) Larval eye pigment cells (orange, arrowheads). B) Wildtype. C) Right and middle eye pigment cells. D) Left eye. E) One right and two left eye pigment cells. Scale bars: 0.5 µm

**Figure 6.**
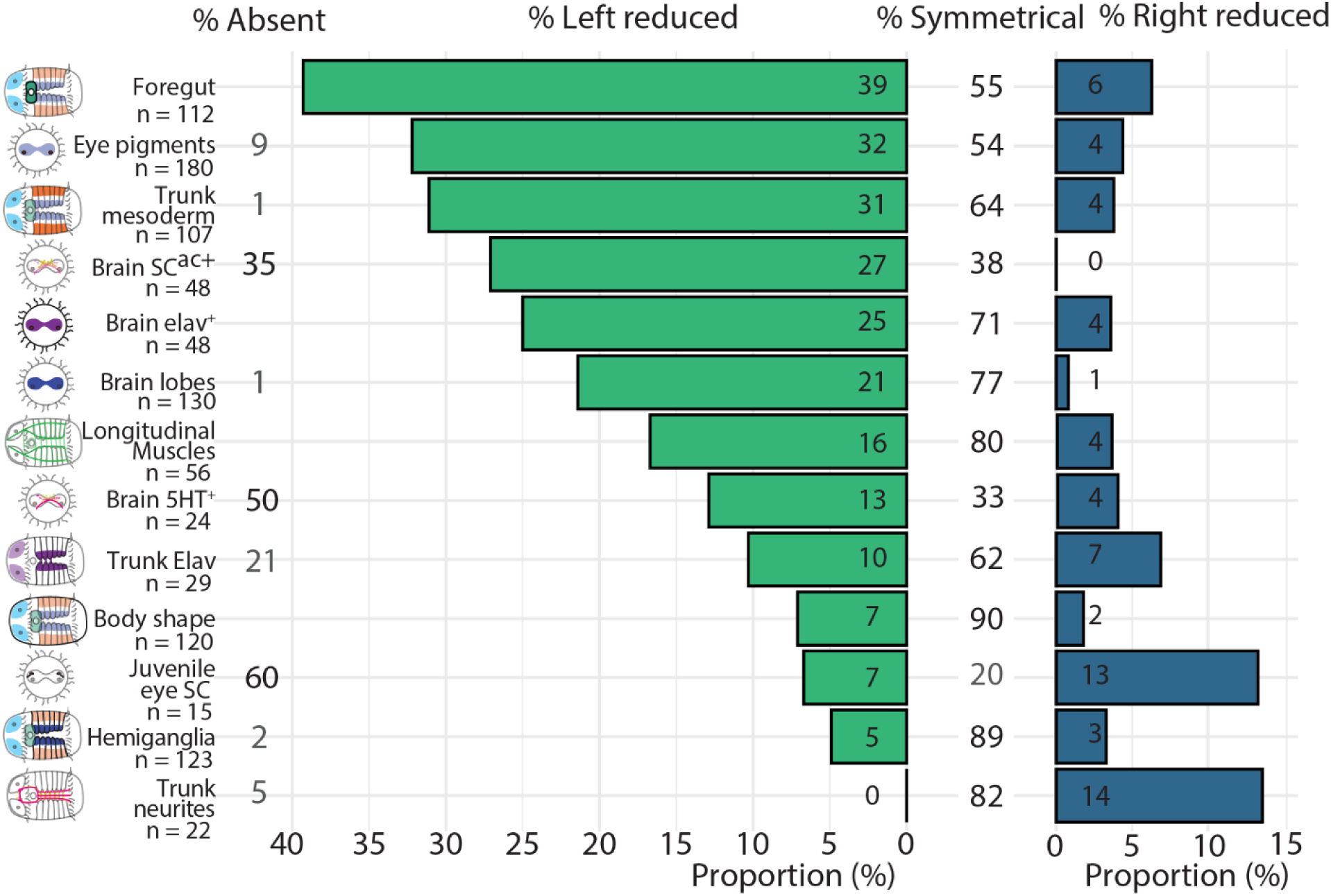
Graphical representation of asymmetries in BMPR1ΔK-injected animals. Asymmetries are sorted from most to least proportion of animals with a reduction of the left side. Percentages are rounded to the nearest whole number.

#### Brain

The majority of BMPR1ΔK-injected animals (n = 130) showed two wildtype brain lobes (concentrations of nuclei in the brain region; 77%; Figs. 4B, 6), with the second most likely phenotype being animals with only a single brain lobe (21%). There was a strong asymmetry in brain lobes; 22% had a reduced or missing left brain lobe, while only 1% showed a reduced right brain lobe (Figs. 4A–D, 6). Brain asymmetry was also apparent in SC^ac+^ cells (n = 48), which were reduced on the left side 27% of the time, and 5HT^+^ cells (n = 24), which were reduced on the left side 12.5% of the time. *Cte-elav1^+^* brain tissue (n = 28) was usually symmetrical (71%), yet a reduction or loss of the left side was common (25%), and a reduction on the right side was rare (4%) (Figs. 4E–L, 6). The level of Cte*-elav1^+^* expression in the episphere was not quantified but was qualitatively slightly lower in BMPR1ΔK-injected animals relative to controls. Notably the majority of animals lacked 5HT^+^ neurons in the brain (50%) compared to only 1% lacking SC^ac+^ cells. In all cases, asymmetric reduction of SC^ac+^, 5HT^+^ or Cte*-elav1^+^* cells was associated with asymmetry in the brain lobes. While the degree of asymmetry varied between measures of brain phenotype, the most likely abnormal phenotype was a reduction or loss of the left brain lobe.

#### Eyes

Approximately 50% of BMPR1ΔK-injected animals (n = 180) had one left and one right orange larval eye pigment cell (wild-type phenotype; Figs. 5, 6; Supplemental Table 1), while 32% of injected animals had a reduction in the number of left eye pigment cells and 4% had a reduction in the number of right eye pigment cells. The majority of these were 1-eyed animals (25% right eye pigment cell only, 2% left eye pigment cell only), but some animals did have medial eye pigment cells or multiple left or right eye pigment cells. 9% of BMPR1ΔK-injected animals had no eye pigment cells. While the number and placement of eye pigment cells varied between abnormal phenotypes, a reduction of loss of left eye pigment cells was most common.

#### Foregut

Foregut tissue (n = 112) in BMPR1ΔK-injected animals showed the greatest degree of asymmetry (Fig. 6), with 39% of animals showing a reduction or loss of the left foregut (Fig. 4D’, arrow denotes the right foregut). 55% of animals show a wildtype, bipartite foregut (Fig. 4B’). Only 6% of animals showed a reduction or loss of the right foregut, and 55% showed a symmetrical foregut. In general, the foregut also appeared slightly smaller than that of uninjected control animals, but this was not quantified (Fig. 4A–D). Foregut asymmetry was the most consistent and striking phenotype showing a loss or reduction of left tissues after BMPR1ΔK-injection.

#### Ventral nerve cord

BMPR1ΔK-injected animals (n = 123) generally showed symmetrical left and right hemiganglia in the VNC (concentrations of nuclei on either side of the ventral midline; 89% Fig. 6), although many hemiganglia appeared farther apart from each other relative to controls (Fig. 4A–D). Very few animals lacked hemiganglia on either side (left-reduced: 5%; right-reduced: 3%). The connectives of the VNC (5HT^+^, n = 22) were generally symmetrical (82%). Of the remaining injected animals, 14% had a loss of the right connectives, none had a loss of the left connectives and 4.5% were lacking connectives altogether (Fig. 4E–H). It is important to note that the first 5HT^+^ connectives originate from neurons on the opposite (contralateral) side of the brain (e.g. 5HT^+^ neurons in the right brain lobe send their axons along the left connectives in the VNC) (38). Cte*-elav1*^+^ tissue in the trunk (n = 29) was usually symmetrical (62%), yet a reduction of either side was common (left-reduced:10%; right-reduced: 7%). Cte*-elav1* expression was not measured but was generally weaker in BMPR1ΔK-injected animals relative to controls (Fig. 4I–L). Strikingly, even in animals with Cte*-elav1* expression in the brain, 21% had no Cte*-elav1* expression in the trunk. Overall, the VNC displayed the least asymmetry after BMPR1ΔK-injection.

#### Mesoderm

In the trunk, BMPR1ΔK-injected animals (n = 107) generally showed mesodermal bands on both sides (64%), but there was a strong asymmetry; 31% had a reduced or missing left mesodermal band, while only 1% showed a reduced or missing right mesodermal band (Figs. 4A–D, 6). This asymmetry corresponded directly to asymmetry in the ventral longitudinal muscle fibers in the trunk (n = 56); all animals with a reduced or missing left or right mesodermal band were also lacking the left or right ventrolateral longitudinal muscle. Of note, only the ventrolateral longitudinal muscles were consistently lost, while the lateral and dorsolateral longitudinal muscles were still intact. Circular muscle fibers were present but were fewer and less organized in regions of the trunk where the anterior-ventral longitudinal muscle was lacking (Fig. 4M–P). In the episphere, the muscle fibers around the brain were not scored because it was unclear whether their asymmetry was due to brain position or mesodermal disruption. Overall non-wildtype individuals showed a clear reduction in the left mesodermal bands that was reflected by a reduction in the left longitudinal muscles.

While most BMPR1ΔK-injected animals (n = 120) were generally straight along their anterior-posterior axis (90%), a small proportion showed a distinct bend in the trunk, mostly concave on the right side (7%) or left side (2%) of the trunk (Figs. 4D’, L’, 6). There was no correlation between body shape and which, if any, other asymmetries were present (e.g. some left and right bent animals lacked the left mesoderm, but not all), and no control animals showed a similar phenotype.

### 3.4 BMPR1dk interacts with BMP4 to change phenotypes

Previously, we reported that *C. teleta* larvae incubated in a 12h pulse of BMP4 protein starting at the 8-cell stage (1q) or just after birth of the 4d micromere (∼64-cell stage, 4q) displayed two different sets of phenotypes, both of which were symmetrical (11). In contrast, when BMPR1ΔK-injected animals were incubated in BMP4 protein for 12h starting at 1q or 4q, they had strikingly different phenotypes than uninjected animals incubated in BMP4 for the same time window. Both 1q and 4q-pulse BMPR1ΔK-injected animals showed asymmetric features, but in opposite directions (Fig. 7).

**Figure 7.**
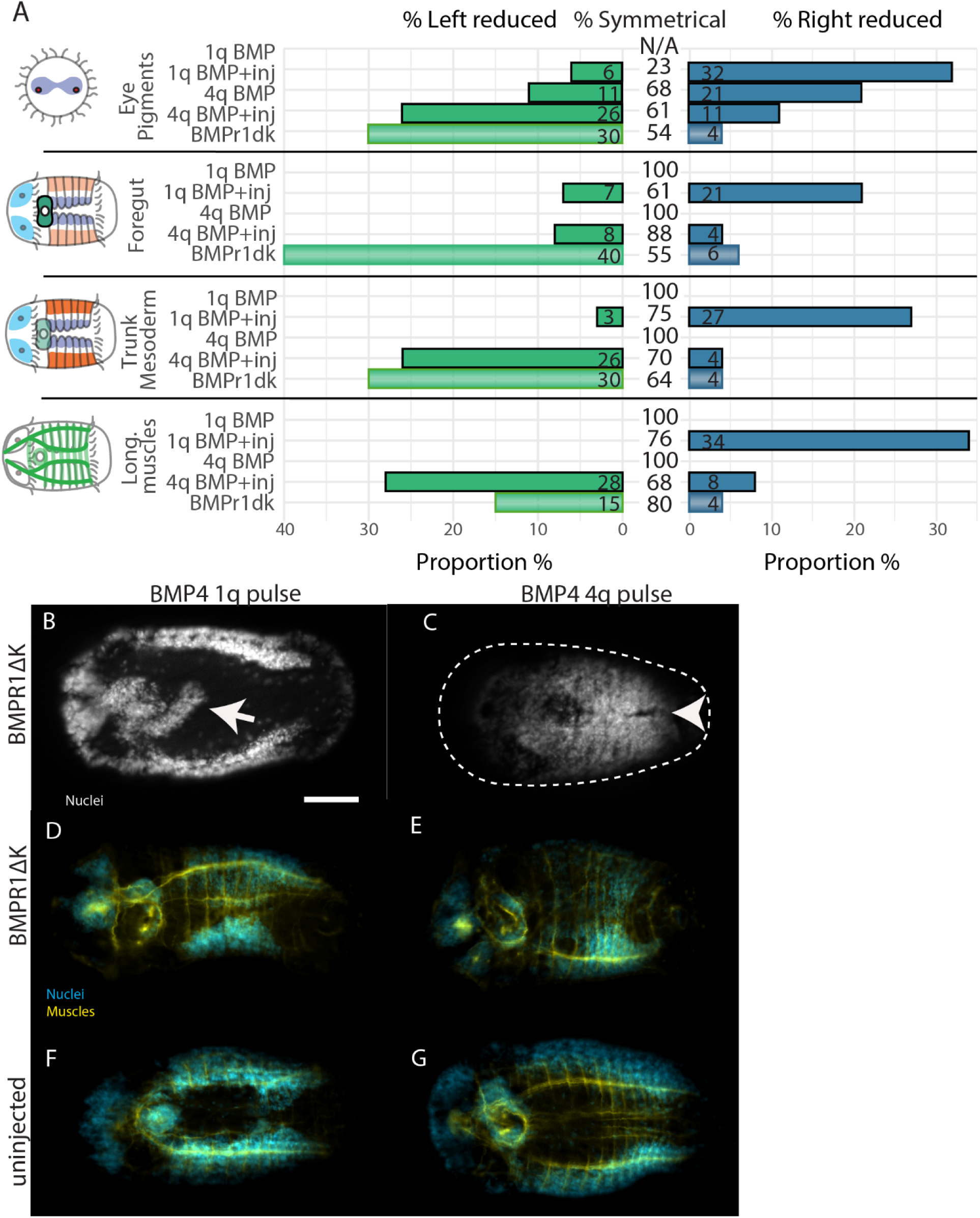
BMPR1ΔK-injected phenotypes after adding ectopic BMP4. A) Graphical representation of asymmetries present in BMP4 + BMPR1ΔK-injected animals. Percentages are rounded to the nearest whole number. B-E) BMPR1ΔK-injected, ventral view. F-G) uninjected, ventral view. B,D,F) 1q BMP 12h pulse. C,E,G) 4q BMP 12h pulse B) BMPR1ΔK-injected + 1q BMP4 animal with elongated foregut (arrow). C) BMPR1ΔK-injected + 4q BMP4 animal with merged hemiganglia (arrowhead). D) BMPR1ΔK-injected + 1q BMP4 animal with reduced right foregut, mesoderm and trunk muscles. E) BMPR1ΔK-injected + 4q BMP4 animal with reduced left mesoderm and trunk muscles. Nuclei: white or cyan; muscles: yellow; scale bar: 50 µm.

#### Eyes – Larval pigments cells

As expected from previous work, uninjected 1q BMP animals had 0 eye pigment cells (n=11) while only 38% of BMPR1ΔK-injected 1q BMP animals (n = 31, Fig. 7) had 0 eye pigment cells. The remainder mostly had 1 eye pigment cell (37%), although some had as many as 3 eye pigment cells (7%). Similarly, while uninjected 4q BMP animals generally had the expected 3 eye pigment cells (n = 15, 87%), only 53% of BMPR1ΔK-injected 4q BMP (n=30) animals had 3 eye pigment cells, the remainder were mostly 2 or 4 eye pigment cells (20% each).

Eye pigment cell asymmetries were quite striking. As presented above, BMPR1ΔK-injected animals had about 30% left-reduced and 4% right reduced eye pigment cell. Surprisingly, BMPR1ΔK-injected 1q BMP animals showed a strong loss of the right eye pigment cell (32%), with only 6% having a loss of left eye pigment cells. In contrast, BMPR1ΔK-injected 4q BMP animals showed a strong reduction in left eye pigment cells (26%), with only a few animals lacking right eye pigment cells (9%). In summary, BMPR1ΔK-injected 4q BMP animals were more similar to BMPR1ΔK-injected animals in lacking left eye pigments, while BMPR1ΔK-injected 1q BMP animals were lacking the right eye pigment cell.

#### Foregut

1q BMP animals generally had a small, symmetrical foregut, similar to previously published results (11). BMPR1ΔK-injected 1q BMP animals (n = 31) also generally had a small foregut, but there was also a general reduction of the right foregut (21%), with almost no loss of the left foregut (7%; Fig. 7). Additionally, 17% of animals showed a posterior elongation from the foregut that was not at all similar to the tripartite foregut seen in 4q BMP animals, but is similar to the right extension of the pharyngeal connection to the esophagus seen later in development (53) (Fig. 7B).

4q BMP animals generally have a tripartite foregut. In contrast, BMPR1ΔK-injected 4q BMP animals (n = 30) did not show any foregut elongation or third lobe, and most had a wildtype foregut. Furthermore, there was no strong asymmetry in the foregut (8% loss of left, 4% loss of right foregut). Broadly speaking, both BMPR1ΔK-injected 1q or 4q BMP animals were more similar to the BMP4 phenotype that the BMPR1ΔK-injected animals which generally has a reduction of the left foregut.

#### Trunk Mesoderm

BMP4 pulses (1q or 4q) generally had no discernable effect on trunk mesoderm. BMPR1ΔK-injected 1q BMP animals (n = 31) had a general reduction of the right trunk mesoderm (27%), and right longitudinal trunk muscles (34%; Fig 7D) with almost no loss of the left mesoderm (3%) or muscles (0%; Fig. 7). In contrast, BMPR1ΔK-injected 4q BMP animals (n = 29) had a general reduction of the left trunk mesoderm (26%; Fig 7E) and left longitudinal trunk muscles (28%) with almost no loss of the right mesoderm (4%) or muscles (7%). Broadly speaking, BMPR1ΔK-injected 4q BMP animals were more similar to BMPR1ΔK-injected animals with a similar loss of the left mesoderm while BMPR1ΔK-injected 1q BMP animals were lacking the right trunk mesoderm.

#### Neural tissue

We previously reported that 1q BMP pulse animals showed a single central brain lobe, while 4q animals had 3 brain lobes, left, right, and midventral. There was no clear pattern in the number or position of brain lobes in BMPR1ΔK-injected 1q or 4q BMP animals. Animals varied from 0–3 brain lobes in various positions. In the trunk, 25% of BMPR1ΔK-injected 4q BMP animals (n = 5/25) had overlapping hemiganglia. This phenotype was reminiscent of the ventral midline loss in animals treated with BMP after 4q (11)(Fig. 7C). In summary, there was too much variation in brain lobe number and location to find a clear pattern in BMPR1ΔK-injected BMP4-treated animals.

### 3.5 Knock-down of *chordin-like* using CRISPR causes right-biased asymmetry

As previously reported, the *C. teleta* genome lacks a homolog of Chordin but has one Chordin-like homolog, *Cte-chd-l*, which is expressed mostly in cleavage-stage embryos and in the foregut, brain, and dorsal midline of developing larvae (47, Webster et al. in prep). In other animals, Chordin-like (Chd-l) is a secreted BMP inhibitor, so we knocked down Cte*-chd-l* using CRISPR to disrupt BMP signaling. Of animals co-injected with Cas9 protein and two gRNAs targeting Cte*-chd-l* (Fig. 8), 21% (10/47) appeared wild-type, and 20% (9/47) failed to gastrulate. Strikingly, a few (9%) lacked the VNC and neurotroch altogether. Most (30%) showed a relatively normal head and small disruptions to the VNC: the VNC connectives were further apart and there were fewer cells in the wider neurotroch (Fig. 8K-O; 14/47). The remainder had a clear asymmetric phenotype, with 22% (10/47) showing asymmetrical features that were weaker on one side compared to the other, as well as a wide, disorganised ventral midline (Fig. 8F-J); 6 had weaker left features, including lack of the left trunk mesoderm (5), a smaller left foregut (3), reduced left hemiganglia (1), fewer left SC^ac+^ cells (Fig. 8F; 2), and fewer left neurotroch cilia (1). Three asymmetrical animals showed weaker right features including a smaller right foregut (2), reduced right hemiganglia (2), fewer right SC^ac+^ cells (1), and fewer right neurotroch cilia (1). Separately, 8 animals showed a separated posterior prototroch region with fewer cilia (Fig. 8N). In summary Cte*-chd-l* CRISPR showed a similar reduction in left trunk mesoderm and foregut compared to BMPR1ΔK-injected animals, but showed very little evidence of affecting the brain.

**Figure 8.**
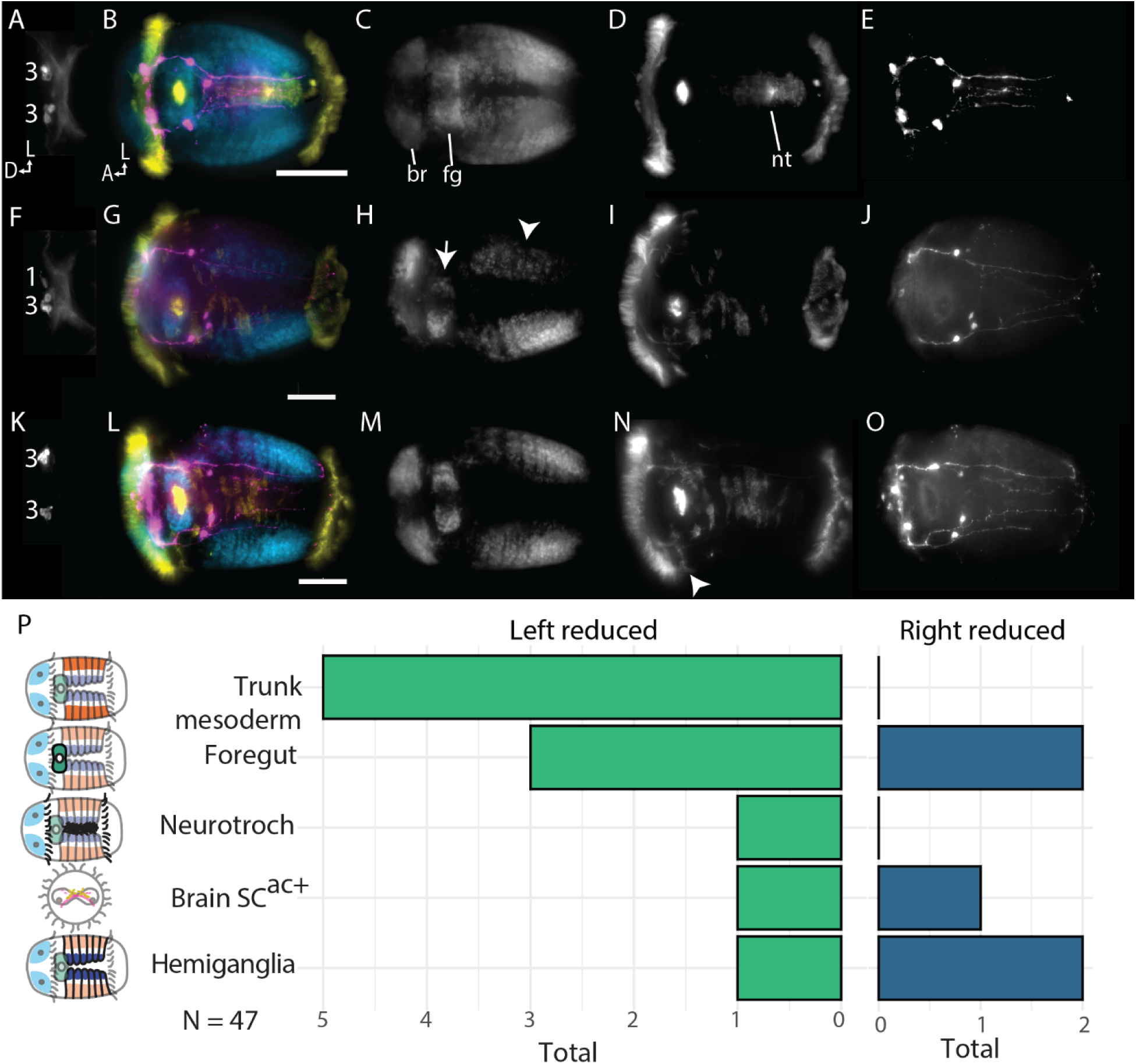
*Cte-Chd-l* CRISPR phenotypes. A-E) Control uninjected animal. F-J) Asymmetrical animal showing loss of left mesoderm (arrowhead) and reduced left foregut (arrow). Note: The apparent asymmetrical brain is an imaging artefact; F is a different animal than G-J. K-O) Animal with wide VNC. P) Summary of asymmetrical phenotypes. A,F,K) Anterior brain showing the number of SC^ac+^ cells. B,G,L) merged images; C,H,M) nuclei (cyan); D,I,N) Acetylated tubulin (yellow); E,J,O) Serotonin (magenta); Scale bars: 50 µm; A: anterior; D: dorsal; L, left lateral; br: brain; fg: foregut; nt: neurotroch.

Supplemental Figure 1. Cte-BMPR1 protein alignment showing the truncation at aa 221 to form Cte-BMPR1ΔK relative to other species. Dme – *Drosophila melanogaster,* Hau – *Helobdella sp*. Austin, Pdu – *Platynereis dumerilii*, Xla – *Xenopus laevis*.

## 4. Discussion

To date, there are only a few papers published that have tested the function of signaling receptors in spiralians; our results add to this work and provide two conclusions that transcend spiralian development. Firstly, we further demonstrate that BMP signaling does not block neural specification or control D-V axis formation in the annelid *C. teleta*, suggesting a lack of conserved function for BMP signaling in these events across Bilateria. Secondly, we report an asymmetrical loss of left tissues in response to a dominant-negative Cte-BMBPR1 construct in *C. teleta*, a phenotype for which we could not find an analog in the literature.

### BMPR1ΔK::mVenus does not increase neural tissue

In vertebrates and insects, opposing gradients of BMPs and their secreted antagonists (e.g., Chordin) generate a gradient of BMP signaling that establishes tissues along the D-V axis. The neuroectoderm is specified in areas with no BMP signaling (e.g. high levels of antagonists), dorsal in vertebrates and ventral in arthropods. If the function of BMP signaling in establishing the neuroectoderm was present in the last common ancestor of Bilateria, we would predict that blocking function of Cte-BMPR1 should expand the domain of neuroectoderm and disrupt or abrogate D-V axis formation (e.g., ventralize animals). In *Xenopus*, a dominant-negative BMP receptor caused a second body axis ventrally (42) and increased neural tissue in animal cap explants (54). In *Drosophila,* injection of dominant-negative forms of BMPR1 orthologs SAX and TKV each caused ventralization of the embryo (loss of the dorsal aminoserosa)(55). We did not observe similar phenotypes in *C. teleta*. Elongated animals showed a relatively normal D-V axis in the trunk, where the expression of *Cte-elav1* in the VNC was generally symmetrically reduced in BMPR1ΔK-injected animals relative to controls. In addition, 5HT^+^ and ac-Tub^+^ cells and neurites were reduced asymmetrically in both the head and trunk of BMPR1ΔK-injected animals. In the brain, neural tissue originating from the right side generally resembled the wild-type condition without evidence of increased neural tissue. Our data further support the hypothesis that BMP signaling is not required for neural delimitation in *C. teleta* and possibly more broadly in Spiralia (10,11,28).

### BMPR1ΔK::mVenus may affect spiralian blastomere quadrant identity

Spiral cleavage is an important aspect of spiralian development, where cell fates can be traced from the earliest cleavages, creating four quadrants, A, B, C and D. In some spiralians, blastomeres in the D quadrant contribute to dorsal tissues while blastomeres in the B quadrant contribute to ventral tissues. Furthermore, in several spiralians, one or more D-quadrant blastomeres acts as the D-V organizer by conditionally specifying fates. This has led to the hypothesis that early during spiral cleavage, the D-to-B axis represents the D-V axis in the larva and in the adult in animals with gradual metamorphosis. In contrast, the A-to-C axis represents right-left axis after gastrulation (25,56). However there is a great deal of unexplored variation in spiralian development (25). Neither *C. teleta* nor *P. dumerilii* follow the D-to-B correlation with the D-V axis (57). For example, in *C. teleta* the left and right trunk mesodermal bands are formed by C- and D-quadrant cells, respectively (49). Fate maps and blastomere isolation studies in spiralians have highlighted how cell fate specification occurs very early in development and can be autonomous, although fates can be labile (58–63). Our previous work suggested that BMP signaling may play a role in establishing quadrant identity during cleavage (11). Specifically, we hypothesized that high levels of BMP signaling (e.g. ectopic BMP) during cleavage shifts blastomeres closer to D-quadrant fates rather than B-quadrant fates. Our data here are less clear. The tissues affected by BMPR1ΔK included tissues mostly from the A-quadrant and D-quadrants: left eye (1a), left-ventral brain (minor contribution from 1a) and left foregut (2a), left-dorsal brain (1d) and left trunk mesoderm (3d). For the A-quadrant, we did not look for an effect of BMPR1ΔK on 3a (head mesoderm) because the episphere was too disorganized, or 4a (endoderm). For the D-quadrant, we did not expect an effect on 2d because it is hypothesized to be specified autonomously and is the organizer, and we did not score an effect on 4d since it makes too few tissues to easily recognize a disruption. Micromeres 1b, 1c, 2c, and 3c did not seem affected by BMPR1ΔK, other micromere fates were not scorable.

If we assume that BMPR1ΔK causes quadrant fate changes in the opposite way to ectopic BMP, we would expect a change from D- to A/C- to B-quadrant fates. We see this for A-quadrant tissue. The lack of left eye and left foregut tissue could be interpreted as a B-quadrant fate. We would also expect D-quadrant tissue to show a C-, B-, or A-like fate. It could be argued that the D-quadrant tissue also shows a B fate, with a lack or reduction of brain tissue and trunk mesoderm. In contrast, we should also expect the C-quadrant to move towards a B-like fate, which we did not find. Instead, the C-quadrant cells appeared to largely produce wild-type tissues with an intact right brain, eye, mesoderm, and foregut. This could indicate that the C-quadrant cells have a separate/redundant, possibly autonomous, mechanism for specification that does not involved BMP signaling. Given these results, we hypothesize that BMPR1ΔK is affecting some of the signals important for quadrant identity, which could explain why A- and D-quadrant cells may be switching to a B-quadrant fate, whereas C-quadrant cells remain relatively unaffected.

Interestingly, the asymmetric phenotype was not the most common phenotype in BMPR1ΔK-injected animals. The majority of all BMPR1ΔK-injected animals failed to develop properly, producing a broad range of unelongated embryos that were difficult to categorize beyond the presence of an anterior-posterior axis. These embryos were more common in higher mRNA injection concentrations and appeared to have a catastrophic disruption of development. The phenotype seen in most elongated, BMPR1ΔK-injected animals may instead represent only a mild disruption of BMP signaling, resulting from some heterogeneity either in terms of concentration, timing, or mosaicism of BMPR1ΔK. Interestingly, adding ectopic BMP did increase the proportion of animals that elongated and more so with a later BMP addition, suggesting some degree of phenotypic rescue (BMPR1ΔK-injected animals: 44% ± 5.3 SE elongation; 1q BMP4 + BMPR1ΔK: 76% elongation, 4q BMP4 + BMPR1ΔK: 88% elongation; 1 biological replicate). Unfortunately, we cannot be more precise because levels of mVenus fluorescence did not correlate with severity of larval phenotypes. These unelongated embryos could represent highly affected animals in which all tissues changed to a B-quadrant fate, leaving little recognizable tissue or axes. It is important to note that B-quadrant cells do not generate “ventral” fates in *C. teleta*, and a shift towards B-quadrant identities would not be a ventralization of the animals.

Given this potential function of BMP signaling in spiralian blastomere quadrant identity, it is possible that D-V axis specification and possibly neural specification was ancestral for bilaterians, but the molecular mechanisms controlling these processes have since diverged significantly in spiralians, especially in Pleistoannelida. Instead, it’s possible that somewhere along the evolution of spiralians, the ancestral gene regulatory network for D-V axis formation (i.e. a BMP signaling gradient combined with activation of the MAPK cascade) shifted to be used for quadrant identity. For example, we previously published that the 5’ end of SMAD1/5/8 mRNA in *C. teleta* is truncated and lacks the nuclear localization signal, something not seen in other spiralians (11). Furthermore, in *C. teleta* Activin/Nodal is required for dorsal-ventral (D-V) axis formation (8) rather than BMP as in leeches (31) and molluscs (28,29). Along this evolutionary trajectory, specification of CNS either diverged so much as to be unrecognizable or came under control of a separate GRN.

### Understanding the molecular effect of BMPR1ΔK::mVenus has a number of confounding factors

The timing of events is critical during development. We showed this previously in the case of BMP signaling during cleavage stages in *C. teleta.* When ectopic BMP was added dramatically changed the resulting phenotype; while adding BMP4 prior to 4q caused a loss of eyes and a radialized brain, adding BMP after the birth of 4d instead caused a third eye and brain lobe to form. While we showed that BMPR1ΔK::mVenus is expressed in injected zygotes, the exact time period of BMPR1ΔK activity could not be determined and may be variable. mVenus fluorescence appeared within a few hours of injection and sometimes lasted for days, and injected embryos without observable mVenus expression showed the same phenotype as animals with mVenus fluorescence. This is not surprising, because dominant-negative mutations can have a strong effect on signaling pathways, even in small concentrations (64). This uncertainty makes it difficult to make hypotheses about the mechanism behind the phenotypes caused by BMPR1ΔK::mVenus because we are lacking data about the timing of BMPR1ΔK::mVenus activity, and early cleavages occur approximately every hour in *C. teleta*.

While almost nothing is known in spiralians, we can make assumptions based on the data available from other animals. Even in well studied groups, the physical interactions between TGF-β family signaling pathways are still being elucidated, making comparisons difficult. For example, Tajer et al. (65) demonstrated that in mice, D-V axis formation requires both BMPR1 and ACVR1, which contradicts our previous understanding of vertebrate D-V axis organization. Not all BMP signaling passes through BMPR1 (ALK3/6), and SMAD1/5/8 is not exclusively phosphorylated by BMPR1. For example, in human myeloma and liver carcinoma cells, Olsen et al (66) showed that ectopic Activin can phosphorylate SMAD1/5/8 via ALK2 (ActivinR1), and knockdown of the type II receptor BMPR2 increases pSMAD1/5/8 via BMP6, BMP7, and BMP9 and ALK2. BMP7 can also phosphorylate SMAD1/5/8 via ALK2 (67). Furthermore, BMPR1ΔK may reduce the availability of BMPR2 receptors in the system due to its effect as a dominant negative in hippocampal progenitor cells (68).

### BMPR1ΔK::mVenus acts early during cleavage

We compared the phenotypes between BMPR1ΔK-injected animals with early ectopic BMP (12h pulse at 1q) to animals with a later ectopic BMP pulse (12h pulse at 4q). In general, the late BMP pulse had little effect on the BMPR1ΔK phenotype, whereas there was a clear difference in the phenotype of early BMP pulse, BMPR1ΔK-injected animals when compared to only BMPR1ΔK-injected animals. Specifically, an early BMP4 pulse reversed the phenotypes, producing a general loss of right tissues instead of left tissues. This fits well with the effects of just a BMP pulse, where the effects of a 1q pulse were dominant to a 4q BMP pulse (11) We predict that the effect of BMPR1ΔK has irreversibly altered fates prior to 4q, otherwise the 4q BMP pulse should have changed the BMPR1ΔK phenotype. A full understanding of the interaction between BMPR1ΔK and BMP requires understanding the timing of BMPR1ΔK activity relative to when specific blastomeres are born. The earliest time we predict that BMPR1ΔK protein is active is around 5 h after injection of the mRNA, or ∼32-cell (3q) stage, when we first see mVenus fluorescence, although there could be a large variation depending on injection timing (± 1 h prior to first cleavage). Thus, BMPR1ΔK activity would begin acting before our 4q BMP treatment window, but after the 1q BMP pulse. This time window of BMPR1ΔK activity suggests that rather than affecting early micromeres directly, it is affecting their more specific daughter cell. For example, BMPR1ΔK likely did not affect 1d specifically to cause loss of the left brain, rather the daughter cells such as 1d^111^. More data will be required to tease apart the complex interactions between developmental timing and the relative timing of BMPR1ΔK and BMP4 activity.

### BMPR1ΔK::mVenus may play a role in left-right axis formation

Our strongest evidence to support the idea that BMPR1ΔK affects left-right specification, whether through quadrant specification or otherwise, comes from the symmetry-reversing effect of BMPR1ΔK injection with an early BMP4 pulse. BMPR1ΔK generally caused a loss of left tissues, whereas adding an early BMP4 pulse caused a general loss of right tissues instead.

In other animals including some spiralians, Nodal and Pitx break symmetry along the left-right axis and disrupting Nodal signaling results in mirror-image symmetry. Nodal is expressed asymmetrically in molluscs (69) and brachiopods (70). Knocking down Nodal signaling in the gastropod *Biomphalaria glabrata* with the drug SB-431542 often disrupted gastrulation, but the other major phenotype was uncoiled shells (69). Adding recombinant Activin also has an asymmetric effect in the snail *Crepidula fornicata* (71). Overexpression resulting in abnormally symmetrical shells and a loss of torsion, presumedly by interfering with endogenous Nodal signaling. These phenotypes, which represent a loss of asymmetry, are in contrast to our reported asymmetrical loss of normally symmetrical tissues. We could not find any other reports of asymmetrical tissue loss due to manipulating signaling pathways for comparison. The closest analog we found is the loss of autonomously-specified tissues after blastomere isolation or ablation, where the cells fated to develop into a specific tissue are removed (58,62,72). For example, in *Xenopus*, while a single blastomere from the two-cell stage will normally form an entire embryo, in the absence of wound healing, that same blastomere will only form a left or right half of an embryo (73).

Preliminary testing of the function of Nodal signaling during *C. teleta* development found no phenotypic effect of adding a wide range of concentrations (25o, 500 and 1000 ng/mL) of recombinant human Nodal protein (R&D Systems 3218-ND) to embryos at different times (data not shown). In addition, ectopic Nodal protein did not affect pSMAD2/3 (D27F4, Cell Signaling) or pSMAD1/5/8 levels during cleavage or gastrulation stages, and ectopic BMP4 protein did not affect nuclear pSMAD2/3 levels (data not shown). It is possible that human recombinant Nodal does not interact with the Nodal signaling pathway in *C. teleta*.

### Chd-l CRISPR affects the trunk and can produce asymmetries

Chordin/Sog is a critical extracellular antagonist of BMP signaling. While many annelids appear to have lost *chordin* from their genomes, they do have a related gene, *chordin-like*, which is thought to act in a similar manner (74). In zebrafish, the Chordin and Chordin-like homologs have at least partially redundant function in regulating D-V axis formation (35), Chd-l transcripts from *Hydra vulgaris* can prevent BMP signaling in zebrafish (75), reinforcing the idea that Chordin-like is also a BMP agonist. *Cte-Chd-l* is expressed throughout larval development, starting as early as 1q (8-cell stage (11)). As a result, we predicted that knocking out *Cte-chd-l* with CRISPR should have a similar effect of adding ectopic BMP4; both should increase the effective availability of BMP in the embryo. This was not wholly the case. While ectopic BMP4 had little effect on trunk morphology except for a loss of the ventral midline and either loss or ectopic formation of foregut tissue, *Cte-chd-l* CRISPR almost entirely affected the trunk, even reducing trunk formation altogether in 9% of cases. *Cte-chd-l* CRISPR also mirrored BMPR1ΔK by producing asymmetric effects. 20% of *Cte-chd-l* Cas9-injected animals had similar asymmetric phenotypes to those found in BMPR1ΔK-injected animals. Unlike BMPR1ΔK-injected animals, phenotypes in the episphere were less common than trunk phenotypes, and there was a lower proportion of animals with a reduction of left tissues.

Trying to decipher how these results inform our understanding of BMP signaling are complicated. We did not examine intermediate developmental stages, so we cannot separate early and late effects of *Cte-Chd-l.* Furthermore, there may be maternal *Cte*-*chd-l* present that could play a critical role early in development and would not be affected by CRISPR. Lastly, not only are the functional differences between Chordin and Chordin-like unclear, but Chordin does not only act as a BMP agonist but can also act as a transporter. In *Drosophila* and *Tribolium* Chordin/Sog binds BMP/DPP and transports it along a diffusion gradient (76), a mechanism that may also affect spiralian development, especially after gastrulation. Overall our *Cte-chd-l* CRISPR results complement the BMPR1ΔK results, especially by producing a similar asymmetric pattern and by having minimal effects on D-V axis and neural tissue.

## Conclusions

Here we have demonstrated the first injected functional mRNA construct in *C. teleta* and identified further questions on the function of BMP signaling. As previously reported, disrupting BMP signaling did not disrupt neural specification. Instead, we think that BMP plays a role in specifying quadrant identity. Specifically, we showed that introducing the kinase-deficient BMPR1ΔK caused a persistent left-reduced phenotype where the left eye, brain lobe, foregut, and mesoderm are reduced or lost, and adding early ectopic BMP4 reversed this phenotype, leading to a similar loss or reduction of right tissues. In addition, we showed that knocking down *Cte-chd-l* with CRISPR/Cas9 gene editing also produced an asymmetric phenotype with a loss of left tissues, but this phenotype was present in the trunk rather than in the episphere. We believe this is a fascinating system to further elucidate the key role that BMPs play in the context of spiral cleavage, and in particular how the rigid framework of spiral cleavage is fundamentally different and can lead to key insights on the general rules of developmental genetics.

## Abbreviations

Alk: Activin Receptor-like Kinase
BMP: Bone Morphogenetic Protein
BMPR1: BMP Receptor 1
BMPR2: BMP Receptor 2
CNS: centralized nervous system
Chd/Sog: Chordin/Short Gastrulation
chd-l: Chordin-like
D-V: dorsal-ventral
SMAD: Suppressor of Mothers against Decapentaplegic
GRN: gene regulatory network, developmental stage: st.
TGF-β: Transforming Growth Factor β.

## Declarations

### Availability of data and materials

The datasets used and/or analyzed during the current study are available from the corresponding author on reasonable request.

### Author’s contributions

The authors confirm contribution to the paper as follows: study conception and design: NBW and NPM; data collection: NBW; analysis and interpretation of results: NBW and NPM; draft manuscript preparation: NBW and NPM. All authors reviewed the results and approved the final version of the manuscript.

### Ethics approval and consent to participate

Not applicable

### Consent for publication

Not Applicable

## Acknowledgements

The authors thank R. Bellin at the College of the Holy Cross for access to their confocal microscope.

Figure captions

See embedded figures.

